# Unveiling functional-metabolic synergy in the healthy brain: multivariate integration of dynamic [^18^F]FDG-PET and resting-state fMRI

**DOI:** 10.1101/2025.05.21.655345

**Authors:** Claudia Tarricone, Giulia Vallini, Giorgia Baron, Erica Silvestri, Tommaso Volpi, Andrei G. Vlassenko, Manu S. Goyal, Alessandra Bertoldo

## Abstract

**Introduction:** Despite accounting for only 2% of body weight, the human brain requires significant amounts of glucose, even at rest, underscoring the importance of functional-metabolic relationships. Previous studies revealed moderate associations between resting-state fMRI functional connectivity (FC) and local metabolism via [^18^F]FDG-PET, yet much remains to be understood, particularly regarding their coupling between functional and metabolic networks.

**Methods:** To this end, we employed multivariate Partial Least Squares Correlation (PLSC) to investigate the functional-metabolic relationship at both nodal and network level. From dynamic [^18^F]FDG-PET data we estimated parameters describing glucose metabolism —delivery rate (*K*1), phosphorylation rate (*k*3), and fractional uptake (*K*i)— and generated within-individual metabolic connectivity (MC) networks. FC was derived from fMRI data filtered into two frequency bands and summarized as region-wise strength to capture nodal characteristics.

**Results:** Our findings revealed that glucose delivery is linked with FC strength, particularly when fMRI signal frequencies include greater hemodynamic contributions. Even stronger functional-metabolic coupling occurs at the network level in the low-frequency fMRI band, with higher MC between sensory/attention and transmodal networks supporting stronger FC within sensory/attention areas.

**Conclusions:** By leveraging PLSC, this work deepens our understanding of the functional-metabolic synergy in the healthy brain, providing new insights into its organization.

**Key Points:**

1. We identified robust functional-metabolic synergy in the healthy brain using multivariate Partial Least Squares Correlation (PLSC), revealing strong associations between fMRI-derived functional connectivity and glucose metabolism from dynamic [¹⁸F]FDG-PET.
2. At the nodal level, glucose delivery (*K*1) is more strongly linked to FC strength than phosphorylation (*k*3) or uptake (*K*i), especially in fMRI bands enriched with hemodynamic components.
3. At the network level, coupling between metabolic and functional connectivity is strongest in the canonical low-frequency fMRI band, suggesting that integrated metabolic support underpins large-scale functional integration across brain systems.

## Introduction

The brain’s remarkable energy demands, consuming approximately 20% of the body’s glucose budget, underscore the critical role of glucose metabolism in sustaining neural function, especially at rest^1^. Understanding the intricate relationship between glucose consumption and functional brain activity is essential for elucidating both normal brain physiology and its dysregulation in disease^2,3^. Recent advances in neuroimaging have enabled the integration of two powerful techniques: [^18^F]-fluorodeoxyglucose positron emission tomography ([^18^F]FDG-PET), which quantifies glucose metabolism, and blood oxygenation level-dependent functional magnetic resonance imaging (BOLD-fMRI), that provides an indirect measure of neuronal activity through hemodynamic changes. BOLD-fMRI has played a crucial role in investigating local brain activity and functional connectivity (FC) at the network level^4–6^. Specifically, investigating how the temporal dynamics of neural and hemodynamic processes in the BOLD signal contribute to its coupling with [^18^F]FDG-PET could reveal novel insights into brain physiology.

Previous studies have examined the relationship between glucose consumption, as measured by [^18^F]FDG-PET, and various fMRI-derived features at both local and global levels^7–12^. Notably, the latest work of Volpi *et al*. (2025) expanded on previous research by analysing not only semi-quantitative measures of glucose metabolism, such as the standardized uptake value ratio (*SUVR*), but also the kinetic parameters derived from Sokoloff’s model^12^. However, while these studies have considered network-level descriptions in terms of FC or graph-derived metrics, they lack a network-level perspective for [^18^F]FDG data. Connectomics has recently shifted its focus towards the molecular processes underlying neural communication, as revealed by PET, with ‘molecular connectivity’ studies^13^. This approach explores the relationships between PET measures across different brain regions (for [^18^F]FDG: metabolic connectivity (MC)). Over the last few years, some studies have begun to investigate MC in relation to FC, but their analyses have been limited to simple correlation or Dice metrics, which cannot capture the nuanced coupling between metabolic and functional networks^14–17^. Additionally, previous studies have considered fMRI signals either with high-pass filtering only (retaining all frequencies above ∼0.008 Hz)^9,11,12,14^ or with band-pass filtering within the 0.1 Hz bands^10,15,18–20^, without accounting for the potential impact of different frequency bands, which may carry distinct biological interpretations and levels of information^21,22^, on their coupling with metabolic counterparts.

To fill these gaps, we propose a novel approach using Partial Least Squares Correlation (PLSC), a multivariate method that enables the investigation of function-metabolic coupling at the individual level. Building upon the work by Volpi *et al.* (2025), we addressed both the nodal coupling between FC strength and [^18^F]FDG kinetic parameters from Sokoloff’s model—specifically the macro-parameter *K*i (tracer’s irreversible uptake) and the microparameters *K*1 (tracer inflow from plasma to tissue) and *k*3 (phosphorylation)—which provide a detailed characterization of glucose metabolism, as well as the network-level coupling between FC and MC.

Additionally, we applied the same analyses across distinct frequency bands of the fMRI signal. These include the canonical low-frequency band (0.008-0.11 Hz), used for BOLD signals to capture low-frequency neural fluctuations^23–26^, and a broader band (0.008-0.21 Hz), which incorporates hemodynamic components^22,27^. While the first frequency band has previously been shown to have a stronger association with metabolic counterparts^25,28^, no study to date has directly investigated functional-metabolic coupling in the second broader band which may better account for potential hemodynamic coupling with [^18^F]FDG metabolism^22,27^.

This analysis allowed us to explore the interplay between neural activity, hemodynamic, and metabolism, offering a more comprehensive understanding of functional-metabolic coupling, at both nodal (*K*1, *k*3, *K*i vs. FC strength) and network (FC vs. MC) level. In addition, we found that multivariate analysis techniques not only revealed robust relationships but also examined patterns of variability across different brain regions and networks, disentangling their different contributions to the covariation between brain function and metabolism.

## Methods

### Participants

Forty-two healthy adults (mean age 58.2±14.5 years, 23 females/19 males) underwent [^18^F]FDG-PET and MRI scans as part of the Adult Metabolism & Brain Resilience (AMBR) study^29^. The study adhered to the principles outlined in the Declaration of Helsinki. Approval for all assessments and imaging procedures was obtained from the Human Research Protection Office and the Radioactive Drug Research Committee at Washington University in St. Louis. Written consent was obtained from each participant. Exclusion criteria for participants included contraindications to MRI, history of mental illness, potential pregnancy, or medication use that could affect brain function.

### Data acquisition

High-resolution structural MRI images were captured using a Siemens Magnetom Prisma^fit^ scanner, employing a 3D sagittal T1-weighted magnetization-prepared 180° radio-frequency pulses and rapid gradient-echo (MPRAGE) multi-echo sequence (TE=1.81, 3.60, 5.39, 7.18 ms, TR=2500 ms, TI=1000 ms, voxel size=0.8×0.8×0.8 mm^3^). The final T1w image was obtained by averaging the first two echoes^30^. Additionally, on the same scanner, T2*w gradient-echo echo planar imaging (GE-EPI) data (TR/TE=800/33 ms, flip angle=52°, voxel size=2.4×2.4×2.4 mm^3^, MBAccFactor=6, a total of 375 volumes acquired over 5 minutes) and two spin-echo (SE) acquisitions (TR/TE=6000/60 ms, FA=90°, with opposite phase encoding directions (AP, PA)) were acquired.

[^18^F]FDG-PET imaging was conducted using a Siemens model 962 ECAT EXACT HR+ PET scanner (Siemens/CTI), following intravenous bolus injection of 5.1±0.3 mCi (187.7±12.1 MBq). PET data were dynamically acquired over a duration of 60 minutes in the eyes-closed waking state. Participant head movements were minimized using a thermoplastic mask. PET data were reconstructed using filtered back-projection (ramp filter, 5 mm FWHM) into 128×128×63 matrices with a voxel size of 2.0×2.0×2.0 mm^3^. The participant’s transmission scan was used to perform the attenuation correction. The reconstruction grid comprised 52 frames of increasing duration, including 24×5 s, 9×20 s, 10×1 min, and 9×5 min frames.

### Structural MRI preprocessing

Structural T1w images were corrected for N4 bias field^31^,skull-stripped^32^, and segmented into grey matter (GM), white matter (WM), and cerebrospinal fluid (CSF) probability maps using Statistical Parametric Mapping tool (SPM12, https://www.fil.ion.ucl.ac.uk/spm/software/spm12). Subsequently, the T1w images were aligned to the MNI152 FSL standard space via nonlinear diffeomorphic registration using the Advanced Normalization Tools (ANTs, v2.4.3)^33^.

The present study employed the cortical parcellation of 74 regions of interest (ROI), clustered from the 100-area Yan homotopic functional atlas^34^, following the procedure presented in Baron *et al.* (2025)^35^ (https://github.com/FairUnipd/ConsensusClustering-YanAtlas). This parcellation is chosen because it can effectively capture the symmetric brain uptake of glucose^36^. Each ROI is assigned to one of the seven Yeo’s resting-state networks (RSNs)^23^: Visual network (VIS) (5 parcels), Somatomotor network (SOM) (13 parcels), Dorsal attention network (DAN) (9 parcels), Salience/Ventral attention network (SALVENT) (10 parcels), Limbic network (Limbic) (5 parcels), Control network (CONT) (13 parcels), and Default mode network (DMN/Default) (19 parcels). Additionally, 12 subcortical and cerebellar regions (AAL2 segmentation^37^) were included: six regions per hemisphere, consisting of thalamus, caudate, putamen, pallidum, cerebellum and hippocampus.

### Functional MRI preprocessing

Functional MRI data preprocessing has been described in previous studies^11,12,17^ and follows an approach similar to the Human Connectome Project minimal preprocessing pipeline^38^. The first four fMRI volumes were discarded to mitigate non-equilibrium magnetization effects. Subsequently, the remaining volumes were corrected for slice timing differences^39^ and magnetic field distortion^40^, then realigned to the median volume using FSL’s *mcflirt*^41^. Next, the FSL’s ICA-AROMA^42^ approach was used for the removal of spurious variance associated with scanner artifacts. A template EPI volume was obtained from realigned fMRI data using *antsBuildTemplate*^33^ and used to estimate an affine transform (*flirt*, FSL) employed to map main T1w tissue segmentations and atlas to the EPI space. To remove spurious sources of variability, motion parameters (rotations, translations), their first-order derivatives, as well as the first five temporal principal components obtained after principal component analysis of WM and CSF EPI signals^43^ were linearly regressed out from all brain voxels in native EPI space^44^.

The temporal traces were high-pass filtered with a cutoff of 0.008 Hz and further low-pass filtered, resulting in two distinct frequency bands of the original BOLD signal: (i) the canonical band between 0.008-0.11 Hz (referred as F1) and (ii) the band between 0.008-0.21 Hz (referred as F2).

### [^18^F]FDG-PET data preprocessing and kinetic modeling

The dynamic PET scans were motion-corrected using FSL’s *mcflirt* algorithm^45^, and to minimize partial volume effects (PVEs), the data were processed without applying any additional spatial smoothing, in agreement with many recent studies^46^. At individual level, PET kinetic modeling was performed on dynamic data as extensively discussed by Volpi *et al*.^17^ (details in the **Supplementary Methods**). Voxel-wise estimation of Sokoloff’s model parameters was conducted with a Variational Bayesian inference framework^47^ to obtain the parametric maps of: blood-to-tissue influx rate constant *K*1 [ml/cm^3^/min], efflux rate constant *k*2 [min^-1^] and the phosphorylation rate constant *k*3 [min^-1^]. Then, the parametric maps of the irreversible tracer uptake, *K*i [ml/cm^3^/min], were obtained from the voxel-level solution of the following equation:

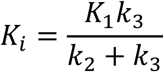

### Within-individual features extraction

For each of the 86 ROIs, pre-processed rs-fMRI time-series, PET time-activity curves (TACs), and the corresponding ROI parametric values were extracted by averaging voxels within the individual GM mask (details in the **Supplementary Methods**). For each participant, two FC matrices—one for each frequency band (F1 and F2)—were obtained by computing Pearson’s correlation coefficients between the time series of every pair of ROIs. To derive a regional measure of functional connection intensity, these matrices were first thresholded at the 80*^th^* percentile, retaining only the top 20% of the correlations^48,49^. Subsequently, the strength of each node was calculated by summing the values of its connections along the rows of the matrix. This process yielded two vectors of node strength (*FCSTR*) for each participant—one for each frequency band (*FCSTR(F1)* and *FCSTR(F2)*). Within-individual MC matrices were computed using the Euclidean Similarity (ES)-based method^17,50^, as briefly outlined in the **Supplementary Methods**.

### PLSC multimodal integration

We applied PLSC to investigate multivariate correlation patterns between various pairs of functional and metabolic measures^51^. Specifically, for both F1 and F2 band-pass filtered FC, we evaluated two levels of multivariate integration: (i) at *nodal-level*, individual regional values of metabolic parameters *K*1, *k*3, *K*i have been related to the corresponding *FCSTR*; (ii) at *network-level*, the edgewise upper triangular features of the full FC and MC matrices have been selected to provide a whole-network assessment of the functional-metabolic coupling. For case (i), the matrices used as input for PLSC were organized as participant-by-metabolic-parameter values on one side and participant-by-*FCSTR* on the other. For case (ii), the input matrices were structured as participant-by-connection. These participant-specific input matrices were then z-scored column-wise (across participants) to ensure a comparable variability range among participants and features. Thus, the resulting multivariate combinations (X-Y) were: *K*1-*FCSTR, k*3-*FCSTR*, *K*i-*FCSTR* for nodal-level investigation, and FC-MC for the network-level. PLSC aims to identify orthogonal latent components by maximizing the covariance between two datasets. Each latent component is associated with two singular vectors, V (i.e., X saliences) and U (i.e., Y saliences), which quantify the contribution of each variable (i.e., ROI or matrix entry) to the functional-metabolic coupling described by that latent component. In the PLSC framework, the linear combinations of the original data variables are weighted by salience maps, resulting in the latent scores. Detailed PLSC equations^51^ are available in the **Supplementary Methods**. **Figure 1** shows the main steps of PLSC analysis, as implemented by the MATLAB *myPLS* toolbox (v1.0) used here^52^, together with the functional and metabolic features considered for nodal or network-level analysis. The statistical significance of multivariate correlation patterns was assessed using permutation testing (200 permutations, critical value of α=0.01)^50,53,54^. The reliability of nonzero salience values was assessed by a bootstrapping approach with 100 random samples, calculating standard scores relative to the bootstrap distributions. The significance of the salience vectors was assessed by checking that their 95% confidence interval excluded zero. Rather, PLSC salience matrices were deemed reliable if their absolute standard score exceeded 3^55^. This threshold was set higher than for salience vectors to account for the larger number of variables^56^.

**Figure 1:**
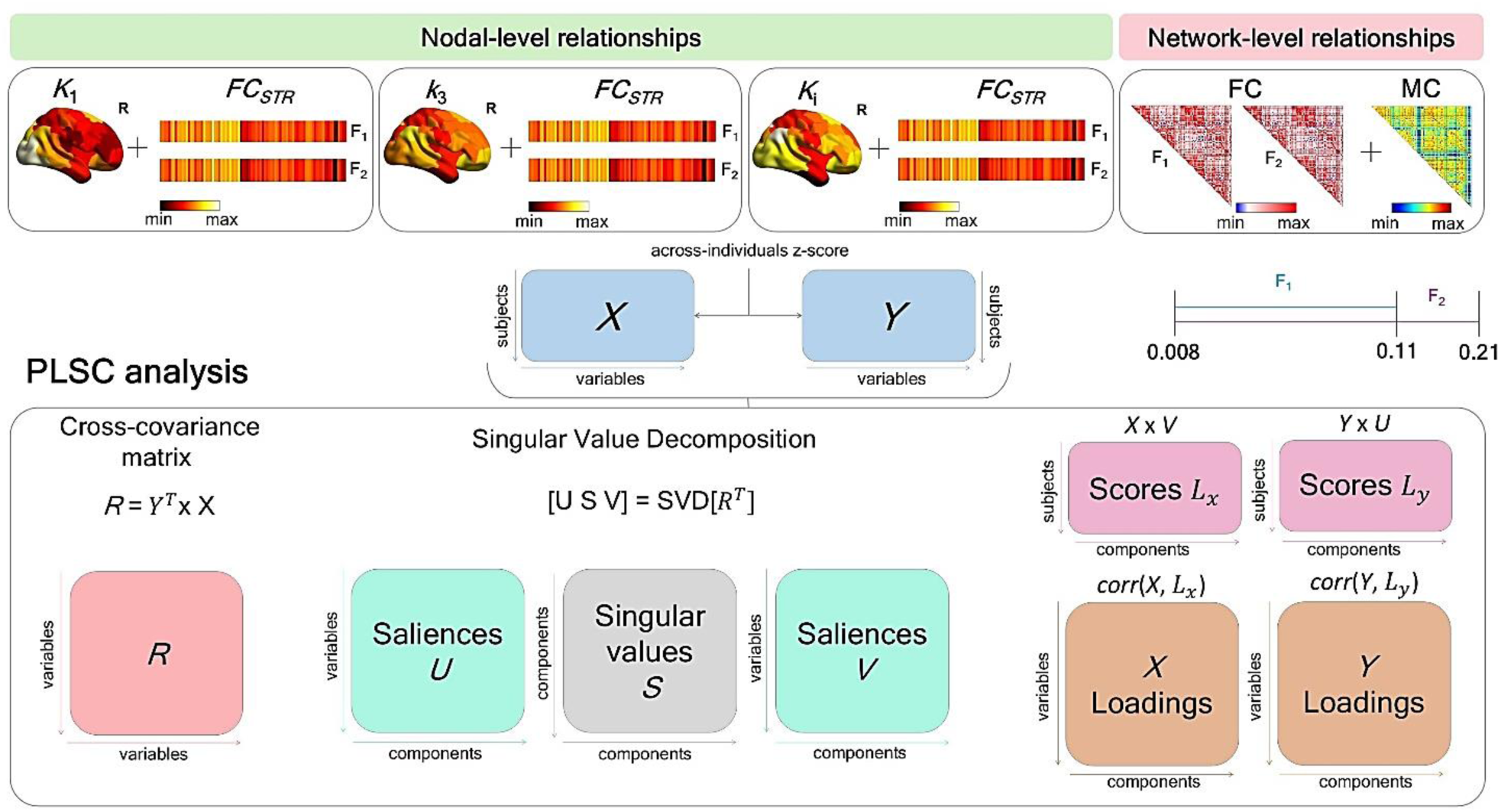
PLSC analysis workflow to assess function-metabolic coupling. At the nodal level, the relationship was evaluated between *K*_1_, *k*_3_, *K*_i_ and FC node strength (*FC_STR_*). At the network level, between the edgewise upper triangles of FC and MC matrices. This workflow was applied twice, considering fMRI features extracted from F_1_ (0.008-0.11 Hz) and F_2_ (0.008-0.21 Hz) frequency bands. PLSC computes the cross-covariance matrix between the original X and Y data matrices, which is then decomposed using singular value decomposition (SVD) into singular vectors *U* and *V* and singular values *S*. This process identifies latent components (LCs) maximizing the shared covariance between the datasets. The PLSC outputs we evaluated include: (i) saliences, i.e. the projections of variables along the identified LCs, which describe their relevance to the coupling, and (ii) scores, representing the projection of each participant’s variable (X or Y) onto the respective saliences pattern (U or V). PLSC also provides X and Y loadings, computed as Pearson’s correlations between the original data (X and Y) and participant-specific scores (*L_x_*, *L_y_*); however, these loadings were not considered in subsequent analyses.

Given the smaller sample size compared to the imaging variables, PLSC models are susceptible to overfitting, resulting in spurious associations that may not generalize well to new data. Therefore, the generalizability of the multivariate correlation patterns obtained from each PLSC analysis was tested using a K-fold cross-validation (7 folds, repeated for 100 iterations), retaining 14% of the participants for the test set in each iteration. After excluding the test participants from the dataset, salience maps were estimated using the training set, and the test-participant data were projected onto these salience maps to obtain the test latent scores^50^. The correlation between the original and test latent scores was then evaluated, and the metabolic-functional pairs were considered generalizable only if the correlation value exceeded 0.5 in at least 50% of the 100 K-fold cross-validation repetitions for both X and Y. Finally, to quantify the relationship between each metabolic-functional pair in the new latent space, Pearson’s correlation between the latent scores (corr(*Lx*,*Ly*)) was computed.

### PLSC output analysis

For generalizable pairs, we conducted a more detailed analysis of the PLSC outputs to assess the contribution of each region/connection to the metabolic-functional coupling. Since the latent scores reflect the alignment between an individual’s original data and the PLSC salience patterns, a positive score indicates a strong similarity between the individual’s data and the salience pattern, suggesting a direct relationship. Conversely, a negative score indicates a mismatch or inverse relationship. Thus, from both a metabolic and functional perspective, participants were grouped based on their individual score values, distinguishing between those with positive and negative scores. Statistically significant differences between these two groups were then assessed through statistical tests both at the nodal and network level. For nodal coupling, we compared the value distributions of each brain region with significant salience between the two score groups using a Wilcoxon rank-sum test (non-normality assessed via Lilliefors test^57^). P-values were Bonferroni-corrected. Then, for each ROI, we calculated the median of the parameter values within each group, computed their difference, and masked the results to retain only significantly changed ROIs. For network coupling, we first computed group-average FC and MC for both score groups, after Fisher z-transforming the individual matrices^58^, and we maintain only edges with significant saliences. We then derived difference matrices by subtracting the negative-score group from the positive-score group for both FC and MC. A Wilcoxon signed-rank test (non-Gaussian distribution confirmed via Lilliefors test^57^) was applied to each network-to-network relationship, with Bonferroni correction for 36 comparisons. Finally, we generated a block-difference matrix by computing the median value within each block (excluding non-significant entries) and retained only those with significant differences. This analysis was performed separately for mean FC (F1 and F2) and MC. The choice of different statistical approaches at the nodal and network levels to examine differences between the two groups of participants based on their score values reflects the different data structures. At the nodal level, we aimed to identify specific ROIs with significant group differences. At the network level, given the matrix-based nature of the data, a difference analysis was more informative for capturing global patterns and network-level changes, rather than edge-wise differences. In addition, testing individual edges would have required extensive correction for multiple comparisons, reducing statistical power. Therefore, a network-based difference approach was statistically more appropriate.

## Results

**Supplementary Figure 1** shows functional (FC matrices and node strength) and metabolic features (MC matrices and glucose micro/macro-parameters) averaged across participants.

### Differences in the FC patterns across frequency bands

The connectograph in **Figure 2a** shows edges where FC differs between the two bands, with red lines indicating stronger FC in F1 than in F2 and blue lines representing the opposite pattern. The thickness of the edges is proportional to the mean FC strength of the connected nodes, and the regions are ordered according to their RSN affiliation, with left-hemisphere nodes appearing before. The distribution of significant differences (|FC(F1)-FC(F2)|>0.05) reveals an overall FC increase in F1 compared to F2 (red edges), particularly within and between networks such as default-mode network (DMN), limbic, salience/ventral attention (SALVENT), and dorsal attention (DAN) networks. In contrast, blue edges, indicating higher FC in F2, are fewer and mainly involve connections within the somatomotor (SOM) and visual (VIS) networks. **Figure 2b** shows the topographic distribution of significant FC strength differences (assessed via Wilcoxon test) between the two frequency bands, with only positive differences (higher *FCSTR* in F1) surviving Bonferroni correction. Increased *FCSTR* in F1 is observed in core DMN regions (posterior cingulate cortex, prefrontal, and temporal areas), the limbic system (orbitofrontal and parahippocampal cortices, temporal pole), and the SALVENT network (insula, medial frontal cortex, parietal operculum). The DAN and SOM networks show significant increases in the postcentral gyrus and motor areas, while subcortical structures (thalamus, caudate, putamen, pallidum, hippocampus) exhibit increased bilateral *FCSTR*. These patterns indicate that frequencies containing more hemodynamic components of the fMRI signal (F2) preferentially support connectivity in unimodal processing regions (more localized within SOM and VIS domains), while lower frequencies (F1) enhance large-scale integration across higher-order cognitive and associative networks.

**Figure 2:**
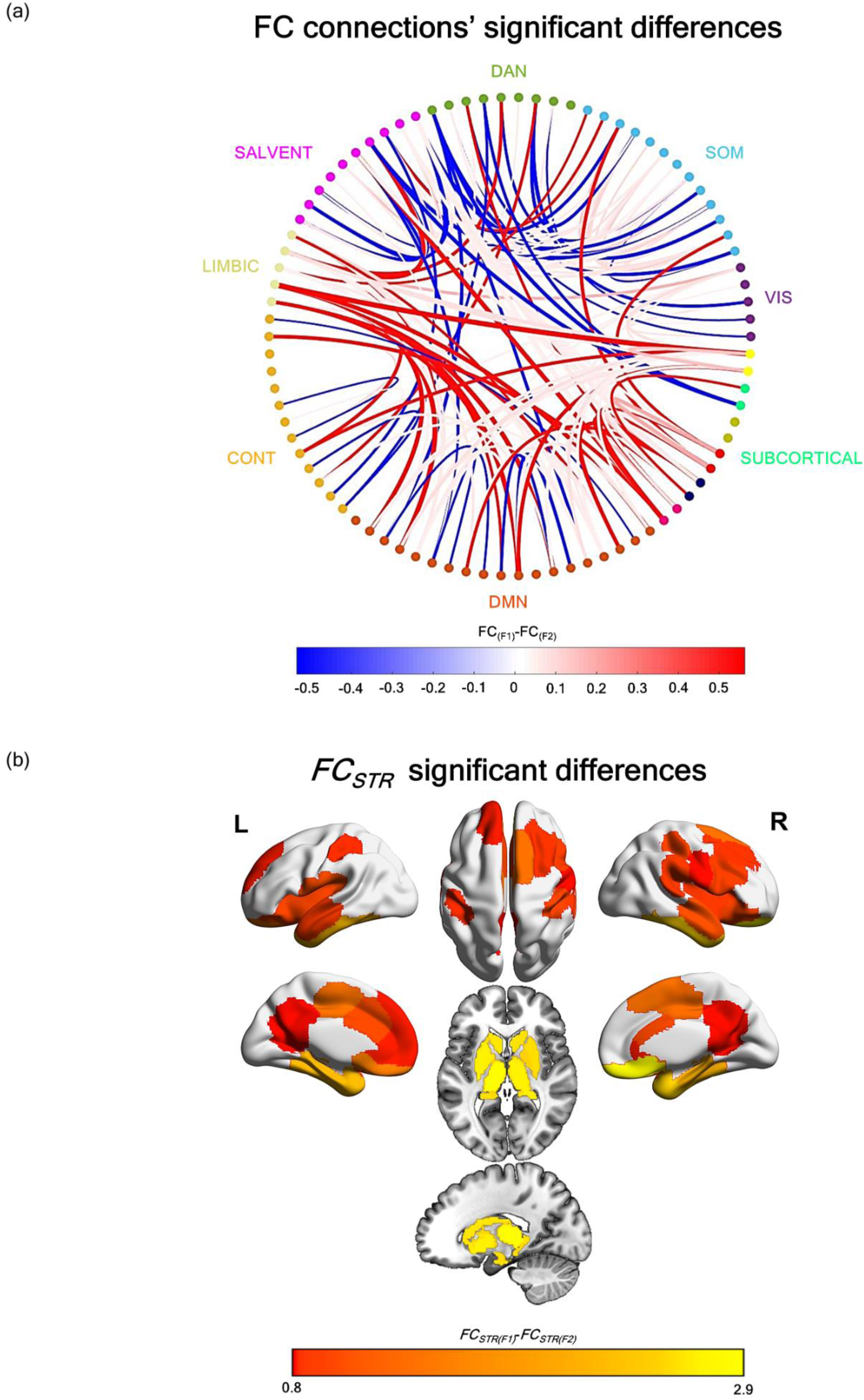
Group-level differences in FC patterns between the two frequency bands. (a) Circular connectivity graph illustrating the significant differences in FC between frequency bands F_1_ and F_2_. FC matrices were first Fisher z-transformed, averaged across participants, and thresholded at the 80^th^ percentile (retaining the top 20% strongest connections). The difference between the thresholded FC matrices was computed, and only connections with an absolute difference greater than 0.05 were retained as significant. Red edges indicate connections where FC in F_1_ is greater than in F_2_, while blue edges represent connections where FC in F_1_ is lower than in F_2_. The thickness of each edge is proportional to the average FC strength of the two connected nodes. ROIs are reordered according to their RSNs (within-network left-to-right hemisphere nodes), which are identified by different colors. (b) Topographic maps of the brain regions (within the 86 clustered ROIs atlas) showing significant differences in FC strength between F_1_ and F_2_, assessed using a Wilcoxon test with Bonferroni correction.

### PLSC multimodal integration

The results of the generalizability test are reported in **Table 1**. For generalizable pairs, the table summarizes the average correlation of all values above the threshold, while for non-generalizable pairs, it includes the average correlation of values below the threshold. Only the *K*1-*FCSTR* (first latent dimension), and FC-MC (first and second latent dimension) pairs were generalizable, while the test failed for *k*3-*FCSTR* and *K*i-*FCSTR*. Specifically, both nodal and network-level pairs were generalizable in F1 and F2 frequency bands, but with different strengths. Indeed, the highest correlation between X-scores and Y-scores (corr(*Lx*,*Ly*)) was found in F2 for *K*1-*FCSTR* pair (r=0.64, p<10^-5^ in F1; r=0.66, p<10^-5^ in F2), while for the FC-MC pair it was observed in F1 second latent dimension (r=0.80, p<0^-9^ in F1; r=0.78, p<10^-9^ in F2). Based on the criteria of the highest corr(*Lx*,*Ly*), in subsequent analyses we considered only the PLSC-identified latent components highlighted in the table. Nodal and network-level PLSC salience maps are shown in **Figure 3**.

**Figure 3:**
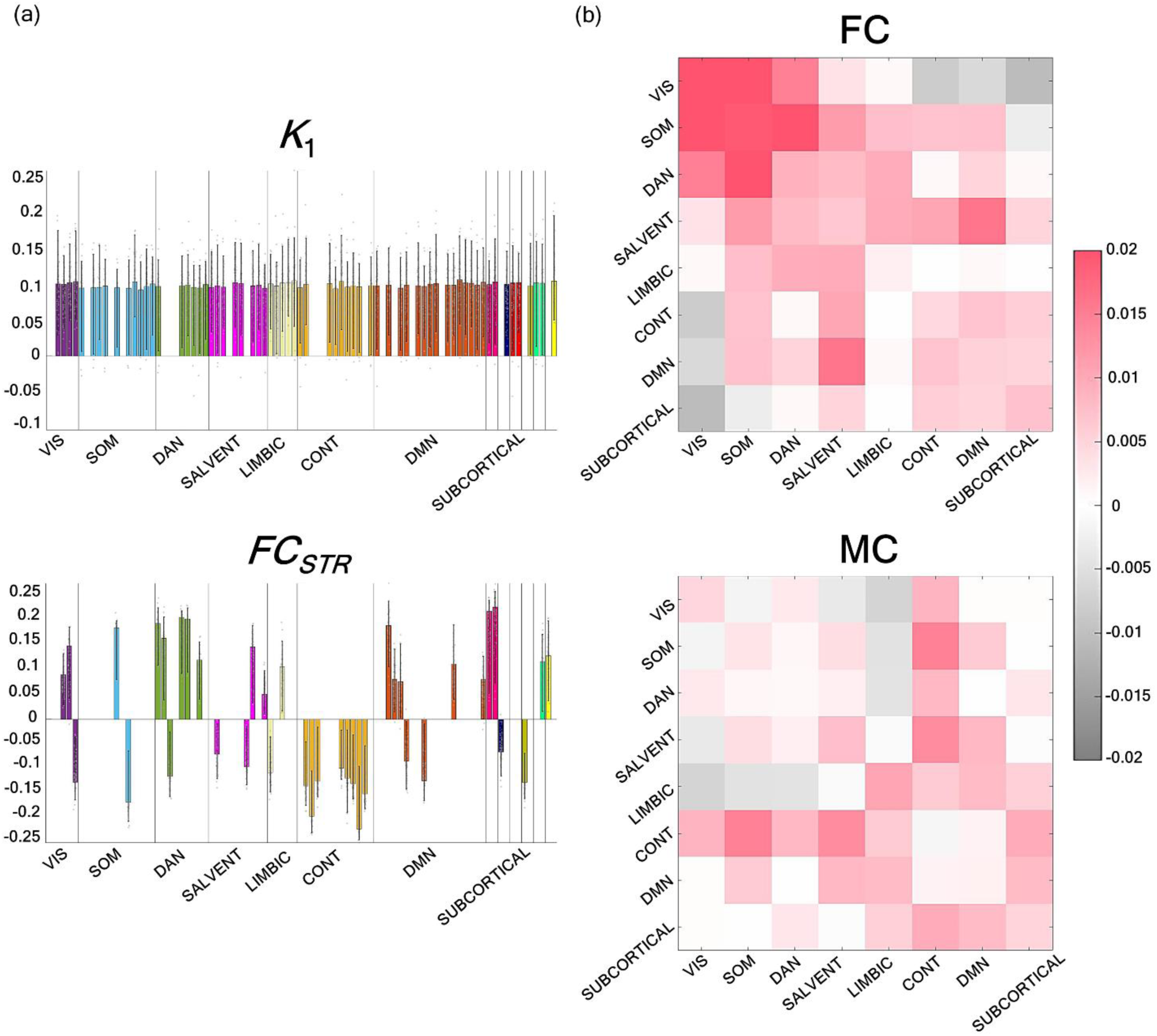
PLSC saliences. a) Nodal-level *K*_1_ and *FC_STR_* PLSC saliences, over the bootstrap samples, showing only the salience weights that are significantly different from zero for each network’s ROIs. RSNs are identified by different colors. b) Network level FC-MC PLSC saliences, reported as the network-by-network average of the statistically significant entries values (treating non-significant saliences as zero, included in the average). While the negative FC saliences between subcortical areas and sensory/attention networks are clearly visible (upper matrix), the same negative entries are less apparent in MC salience pattern (lower matrix) due to the high number of non-significant entries—assumed to be zero, influencing the average—that characterize this interaction

**Table 1:**
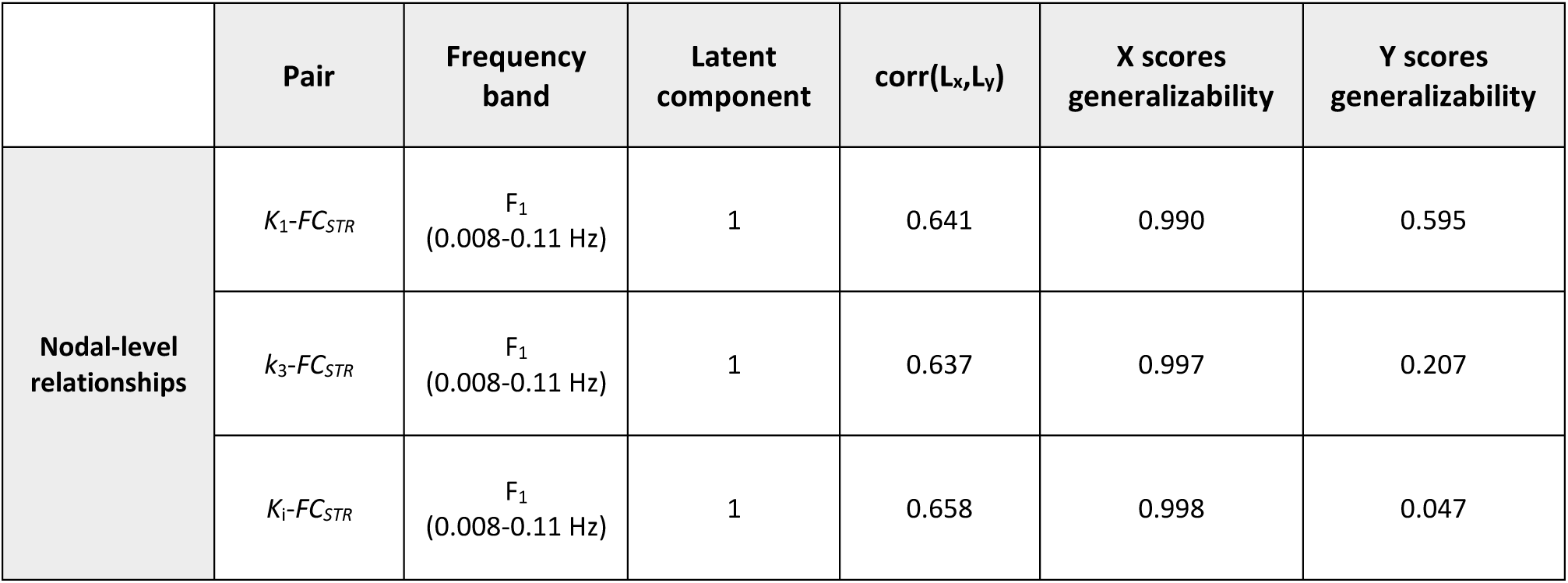
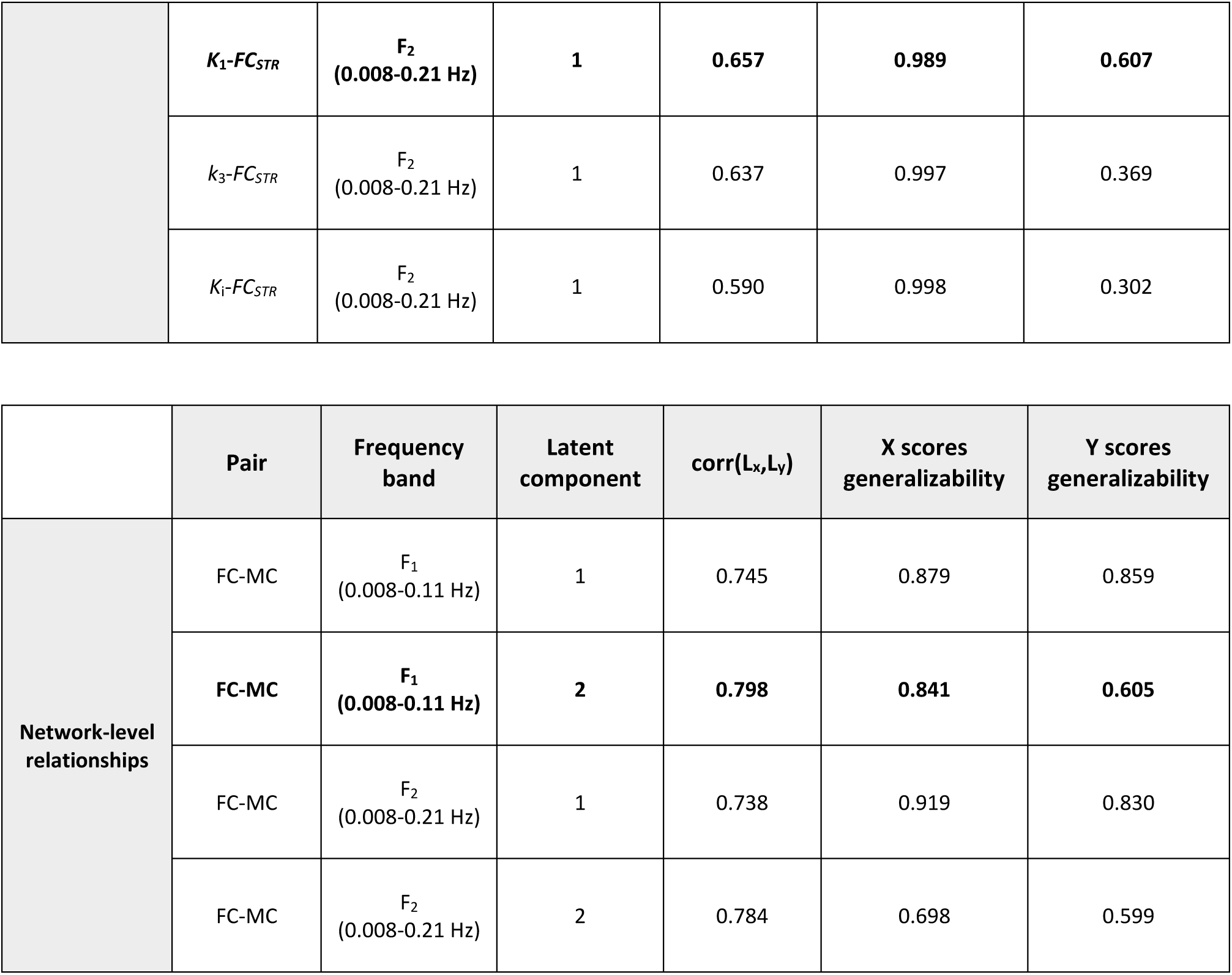
Results of the generalizability test on the PLSC-identified relationships. The upper table reports the results for the nodal-level analyses, and the lower table contains the results for the network-level analysis. For each coupling pair, the ordinal number of the significant latent component (LC) derived from the PLSC analysis and the Pearson’s correlation value between latent score (corr(L_x_,L_y_)), are reported (p-values<0.05 in all cases). The outcomes of the generalizability tests for X and Y are provided in the last two columns. PLSC results are considered generalizable if the correlation between original and test latent scores exceeds 0.5 for both X and Y.

### Nodal-level functional-metabolic relationships: K*_1_-*FC_STR_

At the nodal level (**Figure 3a**), *K*1 saliences (left panel) are uniformly positive and consistent across networks, with relatively homogeneous values. Conversely, *FCSTR* saliences (right panel) exhibit positive values in some sensory/attention regions (i.e. VIS and DAN network), DMN network, and some subcortical regions (especially the thalamus), while negative values prevail in the control network. Consequently, *K*1 and *FCSTR* saliences align in visual, DAN and DMN networks, while also in some subcortical regions, but diverge in the control network. The salience pattern plays a key role in understanding the coupling characteristics in each individual, as the latent scores reflect the alignment between an individual’s original data and the PLSC salience patterns. Therefore, a positive score indicates a strong similarity between the individual’s data and the salience pattern. Conversely, a negative score indicates a mismatch or inverse relationship, where the individual’s data are more closely aligned with the opposite of the salience pattern.

The scatter plot of *K*1-*FCSTR* latent scores in **Figure 4** suggests that two categories of individuals mainly contribute to the variance of the component: one group (in blue) with negative latent scores for both *K*1 and *FCSTR* (i.e., individual’s data more closely aligned with the opposite of the salience pattern) and another group (in yellow) with both *K*1 and *FCSTR* positive latent scores (i.e., strong similarity between the individual’s data and the salience pattern). By masking individual *K*1 and *FCSTR* vectors with their significant saliences, group maps for the two parameters were derived as the median of the values distributions in the respective score groups (*K*1*(POS), K*1*(NEG) and FCSTR(POS), FCSTR(NEG)*). Since positive scores reflect a strong alignment between the individual data and the positive *K*1 salience patterns, *K1(POS)* individuals exhibit data that strongly matches these patterns, i.e. high *K*1 values, whereas *K1(NEG)* individuals show weaker alignment, resulting in lower *K*1 values. Conversely, *FCSTR(*POS) values are higher than *FCSTR(*NEG) in all regions except the control network. The Wilcoxon rank sum test revealed significant differences between the two groups only for *K*1 values, which were higher in the positive group. No significant differences were observed for *FCSTR* values.

**Figure 4:**
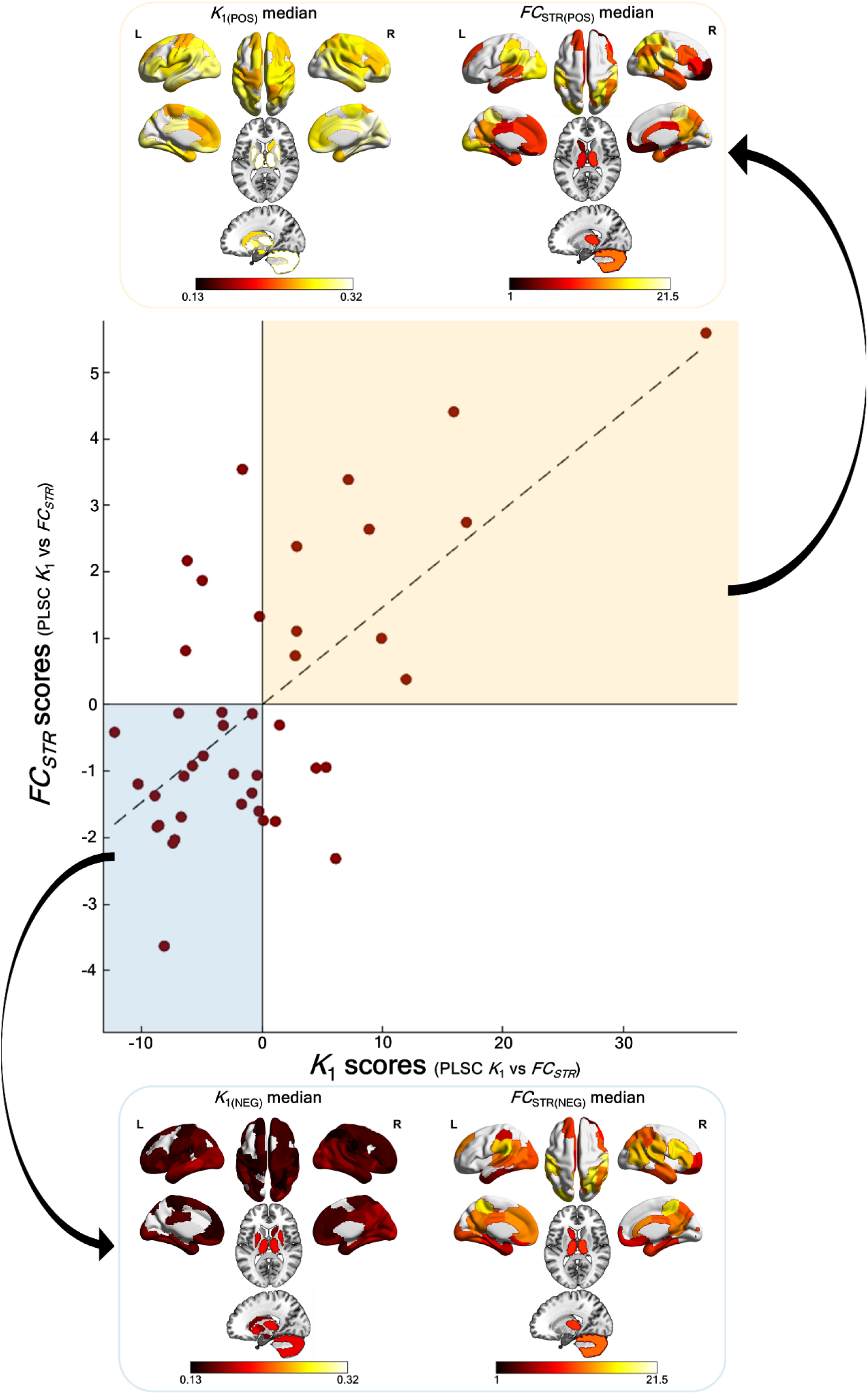
Evaluation of *K*_1_-*FC_STR_* patterns based on the nodal-level PLSC scores model. (*Central panel*) Scatter plot of *K*_1_ vs *FC_STR_* PLSC scores, with each dot representing a single participant. The dashed line indicates the least-square best-fitting line. The highlighted regions (yellow for positive scores, blue for negative scores) identify the participants who primarily contribute to the variance of the PLSC component, exhibiting distinct metabolic-functional characteristics. For each group, the median maps of *K*_1_ and *FC_STR_* are visualized, including only the significant values according to the corresponding PLSC salience map. (*Lower panel*) Negative group; (*Upper panel*) Positive group.

### Network-level functional-metabolic relationships: FC-MC

For the FC–MC pair (**Figure 3b**), we report the network FC and MC salience matrices, i.e. the average of entries in each block (including zeros, i.e. non-significant entries), while full matrices are shown in the **Supplementary Figure S.2**. The FC salience map reveals that sensory/attention networks (visual, somatomotor, and DAN networks) predominantly exhibit intra-network positive weights, while transmodal areas display higher weights in inter-networks connections, especially between salience/ventral attention and DMN networks. Conversely, negative PLSC saliences are more frequent between visual and transmodal networks. In addition, there is a notable reversal in PLSC salience patterns involving subcortical networks, which show predominantly negative weights in the link with sensory/attention networks (except the hippocampus) and positive weights with transmodal networks. The coupling with the metabolic counterpart, as indicated by the MC PLSC-salience map, is characterized by stronger positive weights between sensory/attention and transmodal networks, while lower positive and more negative weights are observed in intra-sensory/attention network connections and their associations with the limbic network. Moreover, subcortical areas exhibit a gradient of connectivity, ranging from negative weights with sensory/attention networks (more clearly visible in the full entries matrix provided in **Supplementary Figure 3**) to positive weights with transmodal networks, supporting findings from the FC PLSC-salience map. In this context, the scores correlation in the latent space is driven primarily by two groups of individuals (**Figure 5**): one with negative FC and MC scores (in blue), showing reversal pattern compared to PLSC saliences, and another with positive FC and MC scores (in yellow), presenting strong alignment between the individual data and the salience patterns. **Figure 5** also reports average matrices for these two groups: FCPOS, FCNEG, MCPOS, and MCNEG. According to the salience pattern, the positive group is predominantly characterized by highly integrated FC patterns within sensory/attention networks and a globally integrated MC, except for the visual and subcortical areas, which exhibit more segregated MC. Conversely, in the negative group, FC patterns showed weaker within-network integration (especially in the visual network) and globally weaker between-network connections, whereas MC showed greater integration within sensory/attention areas. Detailed differences are evident in the full-entry matrices (**Supplementary Figure S.4**), where only the FC mean matrix of the negative group displayed negative relationships, such as anticorrelations between SALVENT and DMN networks, as well as between the limbic and sensory/attention networks (also visible in the block matrix in **Figure 5**).

**Figure 5:**
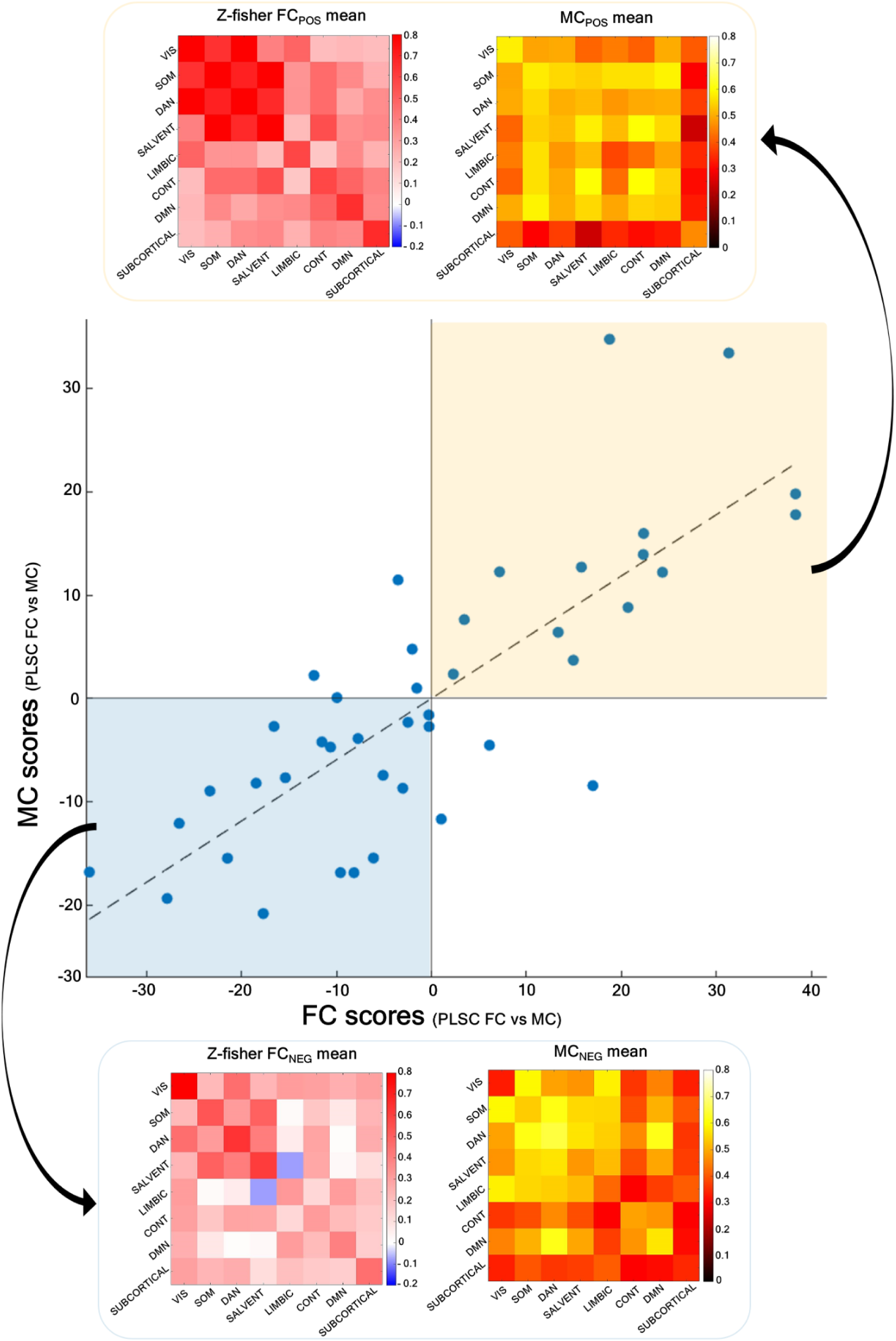
Evaluation of FC-MC patterns based on the network-level PLSC scores model. (*Central panel*) Scatter plot of FC vs MC PLSC scores, with each dot representing a single participant. The dashed line indicates the least-square best-fitting line. The highlighted regions (yellow for positive scores, blue for negative scores) identify the participants who primarily contribute to the variance of the PLSC component, exhibiting distinct metabolic-functional characteristics. For each group, the mean FC and MC matrices are visualized, including only the significant FC and MC entries according to the corresponding PLSC salience map, and the network matrices are obtained by averaging (including zeros) the remaining entries in each block (after Fisher z-transforming the individual matrices). (*Lower panel*) Negative group; (*Upper panel*) Positive group.

The results of the Wilcoxon test between the two groups, shown in **Figure 6**, indicate that the positive group is characterized by significantly higher FC values, especially in the integration between sensory/attention networks and between salience/ventral attention and all other networks (except the visual one), while lower FC values between visual and control networks. This pattern corresponds to significantly higher MC values in the inter-network relations of the control network, especially with the somatomotor and the salience/ventral attention, and to a lesser extent between control, limbic, and DMN networks, while lower MC between DAN and DMN networks.

**Figure 6:**
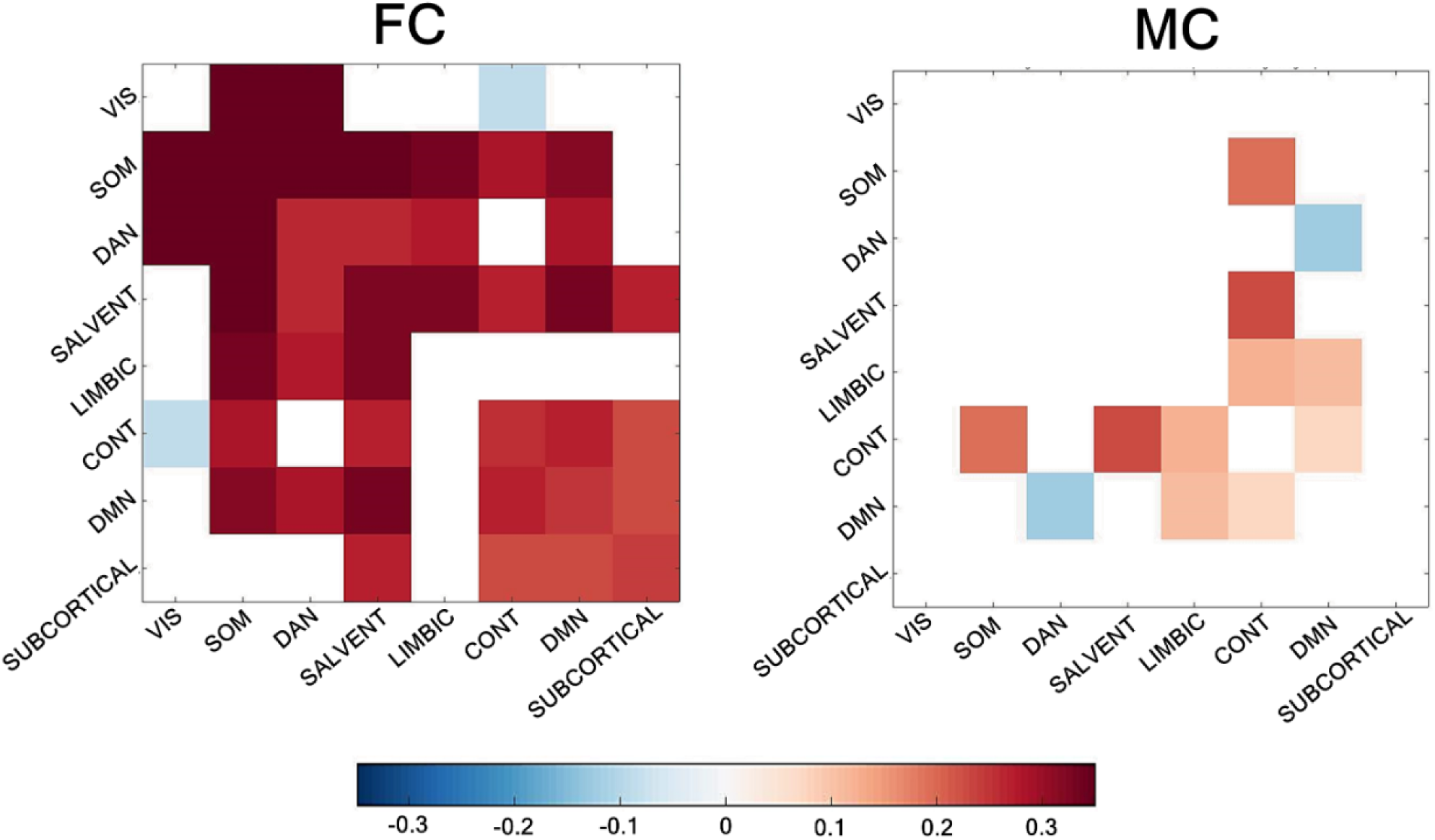
Significant FC and MC inter-group differences. The figure displays significant differences in functional connectivity (FC, left) and metabolic connectivity (MC, right) computed by subtracting group-average values (FC_POS_-FC_NEG,_ MC_POS_-MC_NEG_), followed by a Wilcoxon signed-rank test and Bonferroni correction. The block-difference matrices were generated by calculating the median value of each difference-matrix block (excluding non-significant values, as defined by the salience map). Only blocks showing statistically significant differences between the two groups are visualized.

## Discussion

In this study, we investigated the relationship between brain function and metabolism using the multivariate PLSC approach, which is well-suited for identifying patterns of maximum covariation between two modalities, such as fMRI and PET, while accounting for inter-individual variability^12^. From dynamic PET data, we estimated [^18^F]FDG parameters via kinetic modeling, disentangling the key initial steps of brain glucose metabolism, including delivery (*K*1) and phosphorylation by hexokinase (*k*3), and the overall fractional irreversible metabolic uptake (*K*i), while we also introduced individual-level metabolic connectivity (MC) matrices generated from dynamic PET TACs^17^. Moreover, we investigated the impact of generating FC from fMRI signals filtered into two frequency bands: F1 (0.008–0.11 Hz) and F2 (0.008–0.21 Hz). We hypothesized that different frequency components of the fMRI signal capture distinct neurophysiological processes, which may differentially relate to metabolic activity. This was particularly relevant for the F2 band, where the BOLD signal exhibits a greater hemodynamic contribution^21,22^, potentially affecting its association with [^18^F]FDG kinetic parameters. To explore this, we examined the functional-metabolic relationship at both the nodal (*K*1, *k*3, *K*i vs. *FCSTR*) and the network (FC vs. MC) level, across the F1 and F2 frequency bands.

### Nodal-level functional-metabolic relationships: K*_1_-*FC_STR_

A nodal coupling emerged between *K*1 and *FCSTR*, suggesting that glucose delivery processes play a more significant role in modulating this relationship with the functional counterpart than either glucose phosphorylation (*k*3) or the tracer’s irreversible uptake (*K*i). This finding is consistent with the results of Volpi *et al.*^12^, who showed that *K*1 is the only [^18^F]FDG parameter to exhibit significant bivariate associations with fMRI features (graph measures) related to the hemodynamic response function (HRF) and static FC. Their findings, which highlighted non-linearities and substantial inter-individual variability in the strength of association between metabolic parameters and static FC features, including FC strength, may also explain the limited generalizability of the *k*3-*FCSTR* and *K*i-*FCSTR* relationships. Here, we found that the *K*1-*FCSTR* relationship is stronger in the F2 band, which captures higher-frequency BOLD fluctuations, thereby incorporating more hemodynamic information. Additionally, the F2 band encompasses the frequency range (0.13-0.17 Hz), most likely associated with low-frequency vascular oscillations (or vasomotion)^21^.

According to recent studies^12,59^, perfusion and BBB permeability, as reflected by *K*1, are crucial contributors to the rs-fMRI signal and may be more closely associated with large-scale FC than with glucose metabolism. Having confirmed our hypothesis that the haemodynamic contribution to the fMRI signal might reveal a stronger relationship with metabolic measures reflecting glucose delivery from blood to tissue, we further explored the spatial patterns of these associations using PLSC outputs. This allowed us to examine whether there were differences in the relationship between function and glucose metabolism, both among brain regions belonging to different functional networks (and thus involved in different domains), and between study participants.

Notably, when looking at the salience patterns in **Figure 4a**, positive *FCstr* weights were observed in key DMN regions, as well as in right hemisphere parcels of the visual network, regions of the DAN network and the thalamus. This likely reflects the role of these regions in information processing and integration^7,60^, which requires increased glucose supply^61^. These findings align with previous research linking higher resting cerebral blood flow (CBF) or FDG *SUVR* to stronger FC in the same regions^7,61–63^, but allowing to disentangle the role of glucose delivery from its utilization. In contrast, regions within the control network exhibit negative *FCstr* salience despite positive *K*1 salience. This could reflect a strategy by which the brain optimizes resource allocation to sustain the functional integrity of the cognitive control network, which relies on efficient, dynamic coordination of distributed regions to maintain flexible and adaptive control over cognitive processes, while minimizing energy expenditure^64^. This may be achieved through more efficient inter-regional communication^44^, rather than an increase in metabolic activity.

Additionally, PLSC generates latent scores that quantify associative effects across the sample by projecting the original data onto the PLSC-salience patterns. The joint analysis of saliences and latent scores facilitates the characterization of individual participant profiles. From the examination of the corresponding *K*1-*FCSTR* latent scores (**Figure 4**), two distinct categories of individuals contributing to the variance of the component can be identified. The first group consists of individuals with positive latent scores, characterized by predominantly elevated *K*1 values (relative to the group mean) across the brain, reflecting increased glucose uptake. In this group, high *FCSTR* values are observed in sensory/attention regions, posterior parcels of the DMN, and the thalamus, while lower *FCSTR* values are seen in prefrontal cortex (PFC) areas of the control network. This pattern suggests a reduced number of strong functional correlations between the control network and the other brain networks, i.e., reduced inter-network interactions, possibly favouring localized intra-network processing. This hypothesis is supported by the thresholded mean FC matrices (**Supplementary Figure 3**), which show enhanced intra-network connectivity in individuals with positive latent scores. Conversely, the second group of individuals (negative latent scores) display predominantly lower *K*1 values compared to the group mean, alongside high *FCSTR* in the control network while lower in the remaining regions, possibly reflecting a stronger control inter-network connectivity. This pattern suggests a shift towards between-network communication and synchronization, highlighting the control network’s crucial role in integrating information from networks responsible for processing exogenous, attention-driven information (such as the DAN) and those mediating internally oriented, self-referential processing (such as the DMN)^65,66^. In this group, lower *K*1 values are observed in the PFC regions, reflecting diminished glucose uptake and a reduced need for localized metabolic activity, likely due to the reduction of the high metabolic demands associated with intra-network information processing. Details on the quantitative assessment of the differences observed between intra- and inter-control network relationships in the FC matrices (**Supplementary Figure S.3**) are provided in the **Supplementary Discussion**.

However, despite the presence of a trend across participants transitioning from one group to another, statistically significant differences are evident only for *K*1 values between the two groups, while no significant differences emerge in terms of *FCSTR*. This reflects the limited variability of *FCSTR* values between the two groups, which measure the strength of regional interactions but do not account for the specific connectivity patterns driving these interactions. These observations emphasize the sensitivity of multivariate approaches like PLSC in capturing fine-grained coupling patterns that may not be detectable with simpler bivariate tests. However, while a clear coupling between *K*1 and *FCSTR* exists, nodal-level analysis is inherently constrained by the use of a surrogate connectivity measure (e.g., *FCSTR*).

### Network-level functional-metabolic relationships: FC-MC

Our analyses revealed that the relationship between FC and MC was stronger compared to the one between *FCSTR* and *K*1, reinforcing the initial hypothesis that glucose metabolism and fMRI-derived FC exhibit robust coupling when examined at the network level rather than through local/nodal metabolic measures alone^11^. Additionally, a more robust FC-MC coupling is observed within the F1 band, consistent with prior evidence associating glucose metabolism to slow-frequency fMRI fluctuations within the 0.063–0.098 Hz band^21,67^. As with the node-level analysis, after confirming the validity of our initial hypotheses, we further explored the PLSC results to characterise the patterns of variability-both across networks and between participants - in the function-metabolism relationship at the network level. The observed network-level salience patterns (**Figure 3b)** highlight a key dissociation: while FC showed strong integration between dorsal attention, salience/ventral networks and the DMN, MC primarily linked control network with somatomotor, dorsal attention and salience/ventral networks, as well as subcortical areas with transmodal networks and DMN with limbic/salience systems. This suggests that functional integration predominantly involves interactions between externally oriented (sensory/attention) and internally oriented (DMN) networks, whereas metabolic support is more tightly coupled to the control network’s integration with sensory/attention regions. These findings align with the broader distinction between “extrinsic” systems— characterized by stable, energetically costly FC^68^—and “intrinsic” transmodal networks, which exhibit faster, more dynamic interactions^60^. Notably, our approach extends recent work by Li *et al*.^10^, which used canonical correlation analysis (CCA) to link FC and regional *SUVR* measures. While they reported network-level associations between functional connectivity and regional metabolism, our MC framework provides a more direct comparison with FC and reveals how distributed metabolic patterns support system-level functional communications. Specifically, we found that the control network appears to play a pivotal role in this context, acting as a metabolic and functional bridge between sensory/attention and transmodal networks—a finding consistent with its established role in modulating attention and cognitive control, while also maintaining the balance between externally-oriented and internally-oriented processes^69–71^. This expands on Li *et al*.’s observations by demonstrating that metabolic coupling, not just static local glucose uptake, is integral to large-scale functional network dynamics.

Stratifying participants by scores’ sign (**Figure 5**) further underscored this relationship: the positive group exhibited highly integrated FC within sensory/attention networks and globally distributed MC (except for visual/subcortical areas, which showed metabolic segregation), whereas the negative group displayed weaker FC (particularly in visual networks) but stronger MC integration within sensory/attention regions. This mirrors reports that higher metabolic demand underlies stable FC in extrinsic networks^11^, while intrinsic networks prioritize dynamic flexibility. Crucially, group differences (**Figure 6**) revealed that reduced FC in sensory/attention networks is related with diminished metabolic integration with transmodal regions, suggesting that sustained functional activity in these areas depends on coordinated metabolic support from transmodal systems^50^. Together, these findings highlight a dual regime of functional-metabolic coupling, with the control network serving as a metabolic orchestrator for attention and integration processes.

### Limitations

Despite the innovative findings presented in this study, it is important to acknowledge that fMRI and PET acquisitions were not simultaneous, which may limit the ability to capture concurrent metabolic and functional changes. Additionally, hemodynamic responses are influenced by several physiological and external factors, including hydration status^72,73^, caffeine intake^74,75^, sleep patterns^76–79^, and the time of day^80,81^ at which the scan is performed. However, the relationship between metabolism and function using such a dataset has been explored in previous studies^11,12,29,50^.

### Conclusions

In summary, our results demonstrate the capability of PLSC to uncover covariation patterns between brain function and metabolism at both nodal and network levels, with notable differences based on the BOLD signal band-pass filtering. Specifically, nodal coupling between *K*1 and FC strength is stronger in the frequency band enriched with hemodynamic information. In contrast, network-level relationships, which were also the strongest among all the tested pairs, are more pronounced in the resting state canonical frequency band, associated with BOLD signal frequencies more reflective of metabolic processes. These findings provide compelling evidence for a robust network-level coupling between brain function and metabolism, highlighting the limitations of nodal-level analyses^9,11^. Notably, nodal-level coupling appears to be more influenced by directionality of signal interactions^50^, while the network level gains a more nuanced understanding of the variability in functional-metabolic relationships. Additionally, deepening our understanding of how metabolic energy supports brain functionality has been proven to be essential for unravelling the complexities of brain disorders. By exploring how these intricate relationships are disrupted in pathological conditions, as exemplified in prior research on gliomas using a comparable framework, we can gain critical insights into the specific brain regions responsible for such alterations, ultimately shedding light on the underlying mechanisms that distinguish healthy from diseased states^50^.

## Data availability

The data that support the findings of this study are available from the corresponding author upon reasonable request. The data are not publicly available due to privacy or ethical restrictions.

## Acknowledgments

The author(s) report receiving the following financial support: data acquisition was supported by the McDonnell Center for Systems Neuroscience and NIH/NIA R01AG053503 (AGV) and R01AG057536 (AGV, MSG). Some of the MRI sequences used were obtained from Massachusetts General Hospital; support for the research, authorship and/or publication of this article was provided by #NEXTGENERATIONEU (NGEU) and funded by the Ministry of University and Research (MUR), National Recovery and Resilience Plan (NRRP), Project MNESYS (PE0000006) - A Multiscale integrated approach to the study of the nervous system in health and disease (DN. 1553 11.10.2022).

## Author’s contributions

Andrei G. Vlassenko and Manu S. Goyal collected the data. Claudia Tarricone, Giulia Vallini, and Alessandra Bertoldo designed the research. Claudia Tarricone, Giulia Vallini, Giorgia Baron, Erica Silvestri and Tommaso Volpi analyzed the data. Claudia Tarricone, Giulia Vallini, Giorgia Baron, Erica Silvestri, Tommaso Volpi, Andrei G. Vlassenko, Manu S. Goyal, and Alessandra Bertoldo interpreted the results. Claudia Tarricone and Giulia Vallini wrote the manuscript. All authors revised the manuscript.

## Supplementary material

Supplementary Information is available for this paper.

