## Supplemental Material for "Unveiling functional-metabolic synergy in the healthy brain: multivariate integration of dynamic [^18^F]FDG-PET and resting-state fMRI"

Alessandra Bertoldo, Ph.D.

Via Gradenigo 6/B

35122 Padova, Italy

### Supplementary Methods

*Functional MRI preprocessing*

Each subject was assigned a binary temporal mask to identify brain volumes corrupted by head motion (Framewise Displacement > 0.4 mm). This mask was utilized to adjust the reliability of temporal frames, censoring the data with high-motion frames and checking that each subject had a minimal number of 200 remaining volumes after censoring[1].

*[$^{18}$F]FDG-PET data preprocessing*

PET kinetic modelling was implemented according to Volpi *et al.*[2]. An image-derived input function (IDIF) was extracted from dynamic PET data using a semi-automatic pipeline[3], including 1) internal carotid arteries segmentation; 2) selection of "hot voxels" within the vessel mask; 3) parametric clustering[4] to derive the raw IDIF; 4) IDIF model fitting; 5) Chen's spillover correction[5] using three late venous samples; 6) delay correction. More details are reported in[2,3].

*Common fMRI and PET preprocessing steps*

Preprocessed rs-fMRI signals, along with the regional parametric maps and TACs from dynamic PET images were extracted for each of the total 86 defined ROIs by averaging the voxel activities within the GM SPM mask (a threshold of 0.8 was applied to the probability maps to generate binary masks) for cortical voxels and the non-cerebrospinal fluid (notCSF) SPM mask (threshold of 0.8) for subcortical and cerebellum voxels. The selected segmentation approach was intentionally conservative, yielding a mean sample time course that effectively mitigates partial volume effects (PVEs) while still retaining an adequate number of voxels.

*Within-individual features extraction*

For each pair of TACs, $z_1$ and $z_2$ , their Euclidean distance is calculated as:

$$d_{z1,z2} = \sqrt{\sum_{i=1}^{T}(z_{i,1} - z_{i,2})^2}$$

where $T$ is the number of time points. An index of the ES is derived by subtracting 1 from the normalized Euclidean distance between each regional TAC pair, which is then divided by the maximum distance between TAC pairs, thereby scaling it within the interval [0, 1]. Because of the skewed distribution of ES values, a Fisher z-transformation was applied, followed by a subsequent rescaling of values within the range [0, 1]. This metric has been proven to reveal a network structure in wi-MC matrices, as expected from brain connectivity studies[2].

*Partial Least Square Correlation (PLSC) equations*

As first, the relationship between the two input matrices is quantified by calculating the dot product of the columns of X and Y, given by:

$$R = Y^T X$$

resulting in the cross-product matrix that represents the correlation between the z-scored columns of the X and Y matrices, indicating the correlation between brain functional and metabolic measures. If the columns of the matrices are only centred but not normalized, the *R* matrix reflects covariance. The cross-product matrix, R, undergoes singular value decomposition (SVD[$R^T$])[6,7], resulting in:

$$U * S * V^T = R$$

The decomposition of $R^T$ yields a set of mutually orthogonal latent components (LCs), with U and V being orthogonal matrices containing left and right singular vectors, respectively, and S being a diagonal matrix of singular values. The number L of latent variables equals the rank of the covariance matrix R, which is the smallest of its dimensions. Each latent variable is associated with: (1) a singular value (on the diagonal of S) indicating how much of the cross-block covariance (R) is explained by that latent variable; (2) a vector of singular values V, representing the X saliences; (3) a vector of singular values U, representing the Y saliences. The first and subsequent LCs account for the greatest and progressively decreasing amounts of the cross-covariance matrix (indicated by the square of the singular value), respectively. The saliences indicate how strongly each variable (ROI or matrix entry) contributes to the functional-metabolic correlation explained by a given latent variable. Projecting (using the dot product) each subject's original data (in X or Y) onto the multivariate salience pattern (in V or U, respectively) yields a set of functional-metabolic latent scores $L_X = X * V$ and $L_Y = Y * U$, measuring the similarity of each individual's brain data with the corresponding salient pattern. Finally, functional and metabolic structure coefficients (or "loadings") can be computed as Pearson's correlations between the original data (X and Y) and subject-specific scores ($L_x$, $L_y$). Loadings indicate the extent of contribution of each variable to the latent variable derived from the PLS analysis. In summary, PLS utilizes latent variables to capture the most significant shared information between two different modalities of brain data by maximizing covariance. **Figure 2.** shows the main steps of PLSC analysis, as implemented by the MATLAB *myPLS* toolbox (v1.0) used in this study[8].

### Supplementary Discussion

Specifically, to provide a meaningful and quantitative assessment of the differences observed between intra- and inter-control network relationships in the matrices shown in **Supplementary Figure S.3**, the distributions of intra- and inter-control FC values were compared via Wilcoxon rank sum test, both within the same score group (intra vs. inter -FC within the positive or negative group) and between the two score groups. Notably, and in line with the results of **Supplementary Figure S.3**, significant differences emerged when comparing intra-control FC values between the two groups (p-value = 0.012) and when comparing intra- and inter-control FC within the positive score group (p-value = $10^{-4}$). This supports our idea that lower $FC_{STR}$ values in control regions compared to the average are primarily due to lower inter-network connections of these regions, which are compensated by higher intra-network FC values. Conversely, no significant differences were observed between intra- and inter-control FC in the negative score group (p-value = 0.089), nor when comparing inter-control FC values between the two groups (p-value = 0.534). This suggests that the higher $FC_{STR}$ values in the negative scores group, relative to the average, are due to a balance between intra- and inter-network connectivity, both of which contribute to the overall higher $FC_{STR}$.

### Supplementary Figures

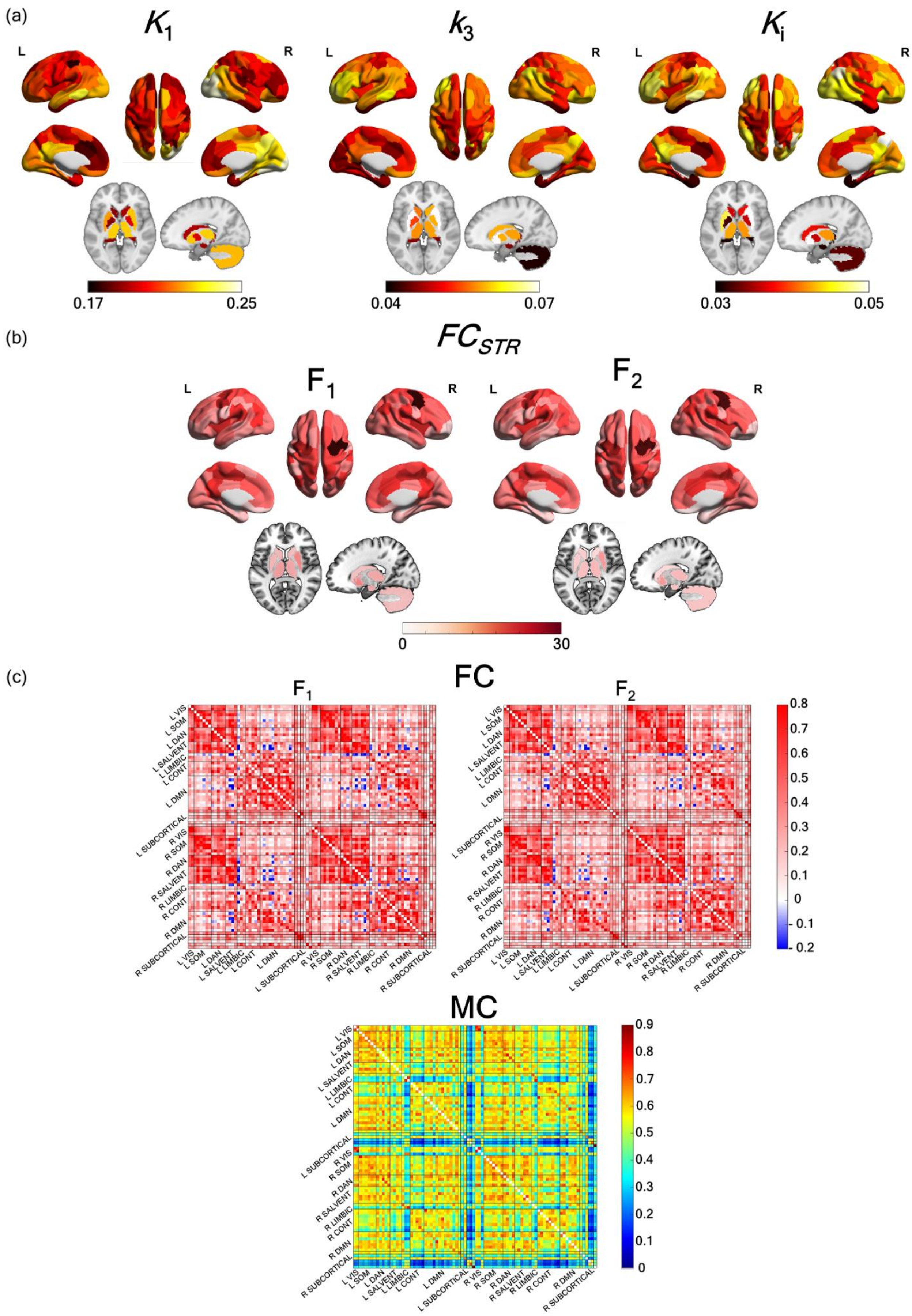

**Supplementary Figure S.1.:** Group-average functional and metabolic features. Dynamic [$^{18}$F]FDG-PET data were used to estimate a) FDG micro/macro-parameters ($K_1$, $k_3$ and $K_i$) at nodal level (86 ROIs) and c) Metabolic Connectivity (MC) matrices at network level (7 RSNs per hemisphere plus 12 SUBCORTICAL regions). From rs-fMRI data, both in $F_1$ and $F_2$ band, we derived c) Functional Connectivity (FC) matrices at network level (7 RSNs per hemisphere plus 12 SUBCORTICAL regions) and b) FC strength ($FC_{STR}$) for each of the 86 ROIs. Brain nodes were grouped based on RSNs (left-hemisphere nodes (L) first, followed by right-hemisphere ones (R)). VIS = visual network; SOM = somatomotor network; DAN= Dorsal attention network; SALVENT = Salience/Ventral attention network; LIMBIC = Limbic network ; CONT = Control network; DMN= Default mode network; SUBCORTICAL = thalamus, caudate, putamen, pallidum, cerebellum and hippocampus.

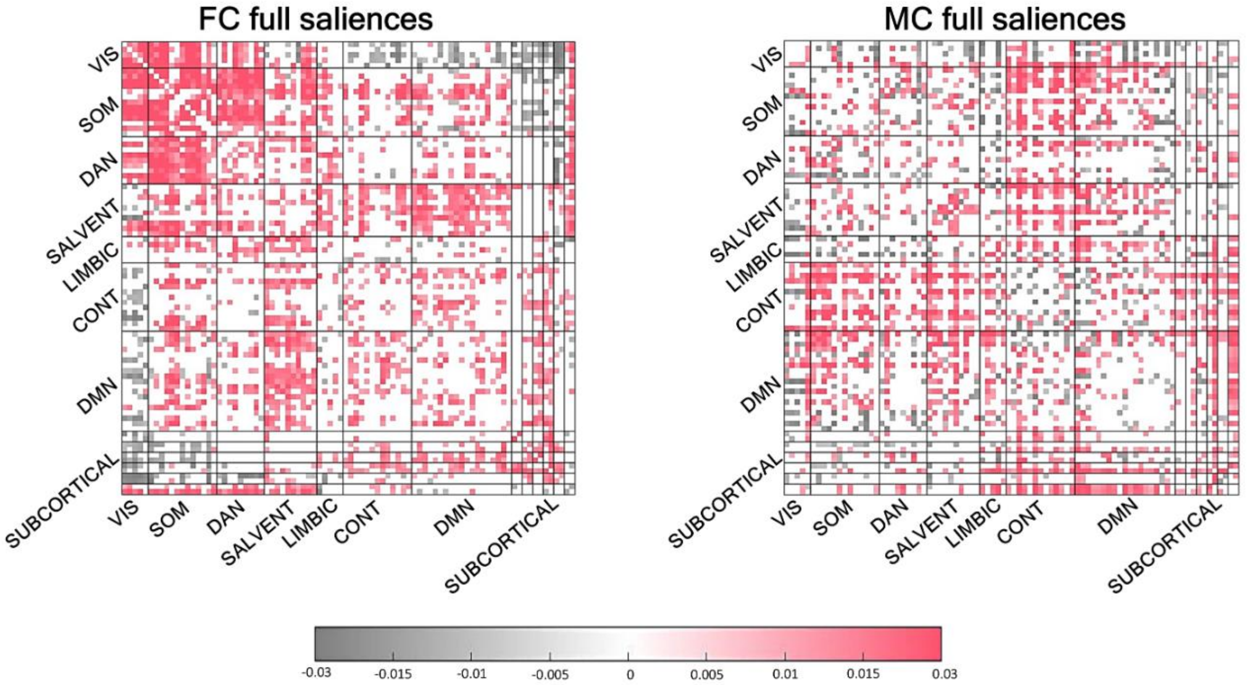


**Supplementary Figure S.2.:** Full matrices of PLSC saliences at network-level. The colours of the entries correspond to the magnitude and sign of the associated significant salience value (i.e., red for positive and grey for negative saliences' values), with non-significant entries in white (according to the bootstrap procedure described in the Methods section).

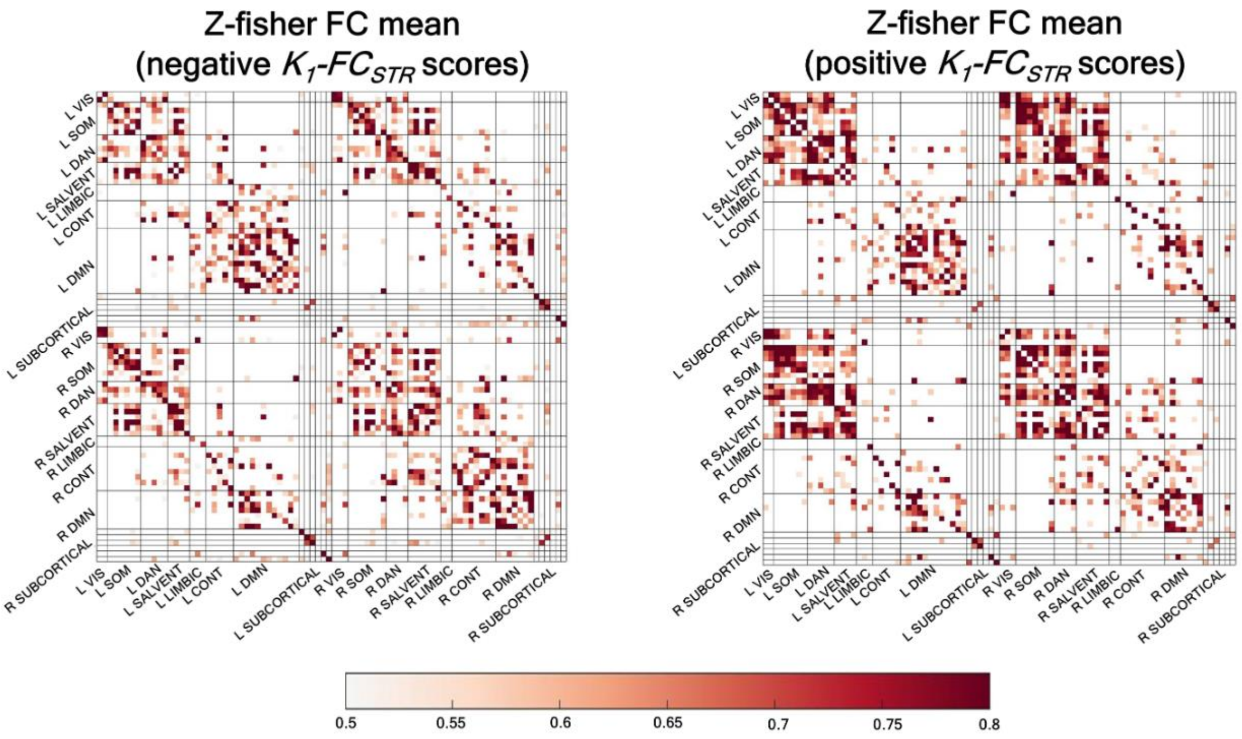


**Supplementary Figure S.3.:** Fisher-z mean and 80th percentile-thresholded FC in the $F_2$ band. The figure displays the mean 80th percentile-thresholded FC matrices for the group of $K_1$-$FC_{STR}$ negative scores' (on the left) and for the group of $K_1$-$FC_{STR}$ positive scores' (on the right), obtained by averaging the FC in the $F_2$ band of the subjects belonging to each of the scores' groups.

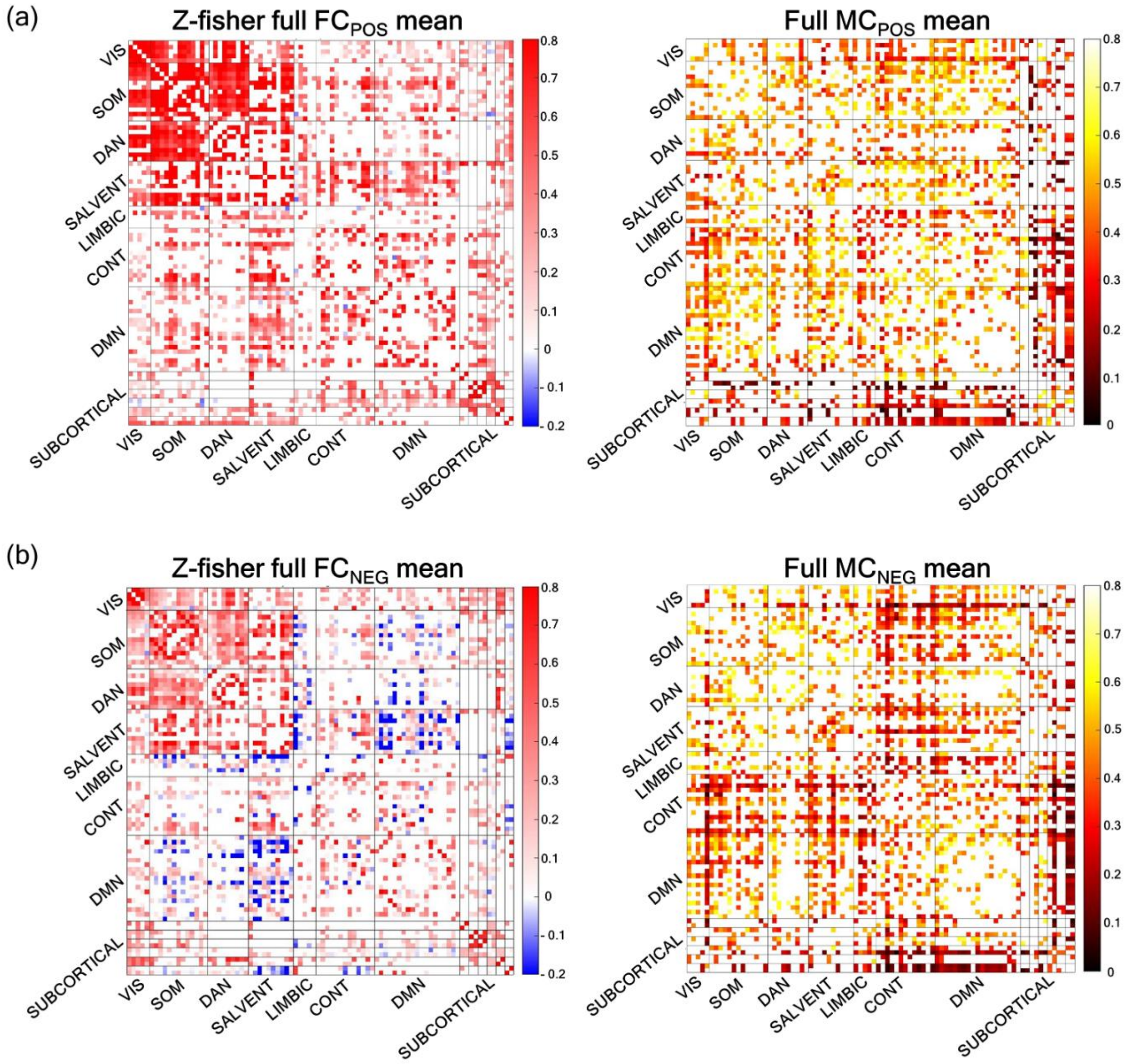


**Supplementary Figure S.4.:** Full-entries scores' groups average FC and MC matrices. a) Mean FC (on the left) and MC (on the right) matrices for the group of subjects with positive FC-MC scores (a) and with negative FC-MC scores (b), obtained including only the significant FC and MC entries according to the corresponding PLSC salience map (non-significant values are shown in white).

8. Zöller D, Kebets V. myPLS - Partial least squares analysis for medical image processing.
